# EffectorK, a comprehensive resource to mine for pathogen effector targets in the Arabidopsis proteome

**DOI:** 10.1101/2019.12.16.878074

**Authors:** Manuel González-Fuente, Sébastien Carrère, Dario Monachello, Benjamin G. Marsella, Anne-Claire Cazalé, Claudine Zischek, Raka M. Mitra, Nathalie Rezé, Ludovic Cottret, M. Shahid Mukhtar, Claire Lurin, Laurent D. Noël, Nemo Peeters

## Abstract

Pathogens deploy effector proteins that interact with host proteins to manipulate the host physiology to the pathogen’s own benefit. However, effectors can also be recognized by host immune proteins leading to the activation of defense responses. Effectors are thus essential components in determining the outcome of plant-pathogen interactions. Despite major efforts to decipher effector functions, our current knowledge on effector biology is scattered and often limited. In this study, we conducted two systematic large-scale yeast two-hybrid screenings to detect interactions between *Arabidopsis thaliana* proteins and effectors from two vascular bacterial pathogens: *Ralstonia pseudosolanacearum* and *Xanthomonas campestris*. We then constructed an interactomic network focused on Arabidopsis and effector proteins from a wide variety of bacterial, oomycete, fungal and animal pathogens. This network contains our experimental data and protein-protein interactions from 2,035 peer-reviewed publications (48,200 Arabidopsis-Arabidopsis and 1,300 Arabidopsis-effector protein interactions). Our results show that effectors from different species interact with both common and specific Arabidopsis targets suggesting dual roles as modulators of generic and adaptive host processes. Network analyses revealed that effector targets, particularly effector hubs and bacterial core effector targets, occupy important positions for network organization as shown by their larger number of protein interactions and centrality. These interactomic data were incorporated in EffectorK, a new graph-oriented knowledge database that allows users to navigate the network, search for homology or find possible paths between host and/or effector proteins. EffectorK is available at www.effectork.org and allows users to submit their own interactomic data.

**Author summary:** Plant pests and diseases caused by bacteria, oomycetes, fungi or animals are threatening food security worldwide. Understanding how these pathogens infect and manipulate the host is key to develop sustainable crop resistance in the long term. Effector proteins are secreted by pathogens to subvert the host immune responses. The roles of several effector proteins have been described; however, it is yet poorly understood how effectors interact with host proteins at a global level. To address this issue, we have generated EffectorK, an interactive database focused on the model plant species *Arabidopsis thaliana*. This database contains manually curated Arabidopsis-effector protein interactions from the available literature on a wide variety of pathogens. It also contains new experimental data on effectors from two vascular pathogens: *Ralstonia pseudosolanacearum* and *Xanthomonas campestris*. This work integrates all the gathered knowledge over the last decades and allows to identify general patterns of how effectors interact with the host proteome. This knowledge is easily accessible and searchable at www.effectork.org.

## Introduction

Plants are continuously confronted with a wide variety of pathogens including bacteria, oomycetes, fungi, nematodes and insects. To prevent their proliferation, plants have evolved a complex multilayered immune system [1]. The first layer of defense corresponds to constitutive physical and chemical barriers such as the cuticle, cell wall or secondary metabolites [2, 3]. Plants are also able to recognize highly conserved pathogen-associated molecular patterns (PAMPs) through pattern-recognition receptors triggering induced defense responses collectively known as ‘PAMP-triggered immunity’ (PTI) [4]. These responses are usually enough to prevent most potential invaders; however, some pathogens secrete effector proteins to subvert the defense responses and alter diverse cellular processes to ease their proliferation [5]. Plants, on the other hand, have evolved several intracellular nucleotide-binding site-leucine-rich repeat (NBS-LRR) receptors recognizing these effectors and activating potent defense responses collectively known as ‘effector-triggered immunity’ (ETI) [6].

Although the targets and molecular functions of some effectors have been well characterized [7–10], for most effectors they are still unknown. The main factors complicating the large-scale identification and characterization of effector-host protein interactions are: the wide diversity of pathosystems, the difficulty to identify *bona fide* effector genes, the collective contribution of effector proteins, the complexity of the host responses and the lack of robust high throughput techniques. For the model species *Arabidopsis thaliana* (*Ath*), to our knowledge, there are only two studies in which systematic effector-host protein interactions at the effectome-scale have been identified [11, 12]. In these studies plant targets of effector proteins from *Pseudomonas syringae* (*Psy*, bacterium), *Hyaloperonospora arabidopsidis* (*Hpa*, oomycete) and *Glovinomyces orontii* (*Gor*, fungus) were identified by yeast two-hybrid (Y2H). They reported that the effectors of these three species converged onto a limited set of *Ath* proteins. These studies also demonstrated that many effector targets are important for plant immunity and showed that their importance correlates with the level of effector convergence.

Bacterial wilt, caused by *Ralstonia pseudosolanacearum* (*Ralstonia solanacearum* phylotype I, *Rps*), and black rot, caused by *Xanthomonas campestris* pathovar *campestris* (*Xcc*) are listed among the top five plant bacterial diseases in the world [13]. Both *Rps* and *Xcc* are xylem-colonizing pathogens and rely on their type III secretion systems for full virulence [14, 15]. This ‘molecular syringe’ allows the pathogen to deliver type III effector proteins (T3Es) directly into the host cell in order to promote disease. The roles of several of their T3Es have been characterized [16, 17], but most knowledge on T3E functions comes from the study of *Psy*, which resides on leaf surfaces and in the leaf apoplast [7, 18]. Focusing mainly on a few species offers a partial view of effector biology. It is therefore crucial to expand our studies to other species to grasp most of the existing diversity of effector proteins and pathogen lifestyles.

To obtain a deeper understanding of the global *Ath*-effector protein interactome, we conducted two systematic large-scale screenings with T3Es from *Rps* and *Xcc*, the first vascular pathogens screened in this manner. Additionally, we conducted an extensive literature survey to gather published *Ath* targets of effector proteins from pathogens from four different kingdoms of life: Bacteria, Chromista, Fungi and Animalia. Combining all these data allowed us to identify 100 new ‘effector hubs’ (i.e., *Ath* proteins interacting with two or more effectors). Together with *Ath*-*Ath* protein interactions retrieved from public databases, we generated a comprehensive *Ath-* effector protein network that captures the wide diversity of *Ath* pathogens. This network allowed us to detect general trends of effector interference with the host proteome. We have created a publicly available interactive knowledge database called EffectorK (for Effector Knowledge) which allows users to access and augment this network.

## Results

### Systematic identification of Arabidopsis targets of *R. pseudosolanacearum* and *X. campestris* effectors

Multiple Y2H screenings were performed to identify *Ath* targets of *Rps* and *Xcc* effector proteins. In a first screening, we identified 42 *Ath* targets for 21 out of 56 T3Es from *Rps* strain GMI100 screened against a library of more than 8,000 full-length *Ath* cDNAs (8K space). In the second and third screenings, we identified 176 *Ath* targets for 32 out of 48 T3Es from *Rps* strain GMI1000 and 52 *Ath* targets for 18 out of 25 T3Es from *Xcc* strain 8004 screened against an extended version of the previous library containing more than 12,000 *Ath* full-length cDNAs (12K space) (S1 Fig and S1 Table). On average, T3Es from *Rps* interacted with 10.7 *Ath* proteins while T3Es from *Xcc* interacted with 5.3 *Ath* proteins. These *Ath* cDNA libraries had been previously used to test interactions with effector proteins from *Hpa*, *Psy* (8K space) and *Gor* (12K space) [11, 12]. The subset of interactions of effectors from *Rps*, *Xcc* and *Gor* in the 8K space was used to compare with previously published *Hpa* and *Psy* data (Fig 1). In general, *Rps* effectors interacted on average with more *Ath* proteins than the other screened species; however, this difference is only statistically significant when compared to *Gor* effectors (one-tailed Wilcoxon signed-rank test p-value = 0.0005). These data show that effector proteins from these five different species, on average, tend to interact with a similar number of *Ath* proteins regardless the kingdom, lifestyle or effectome size.

**Fig 1.**
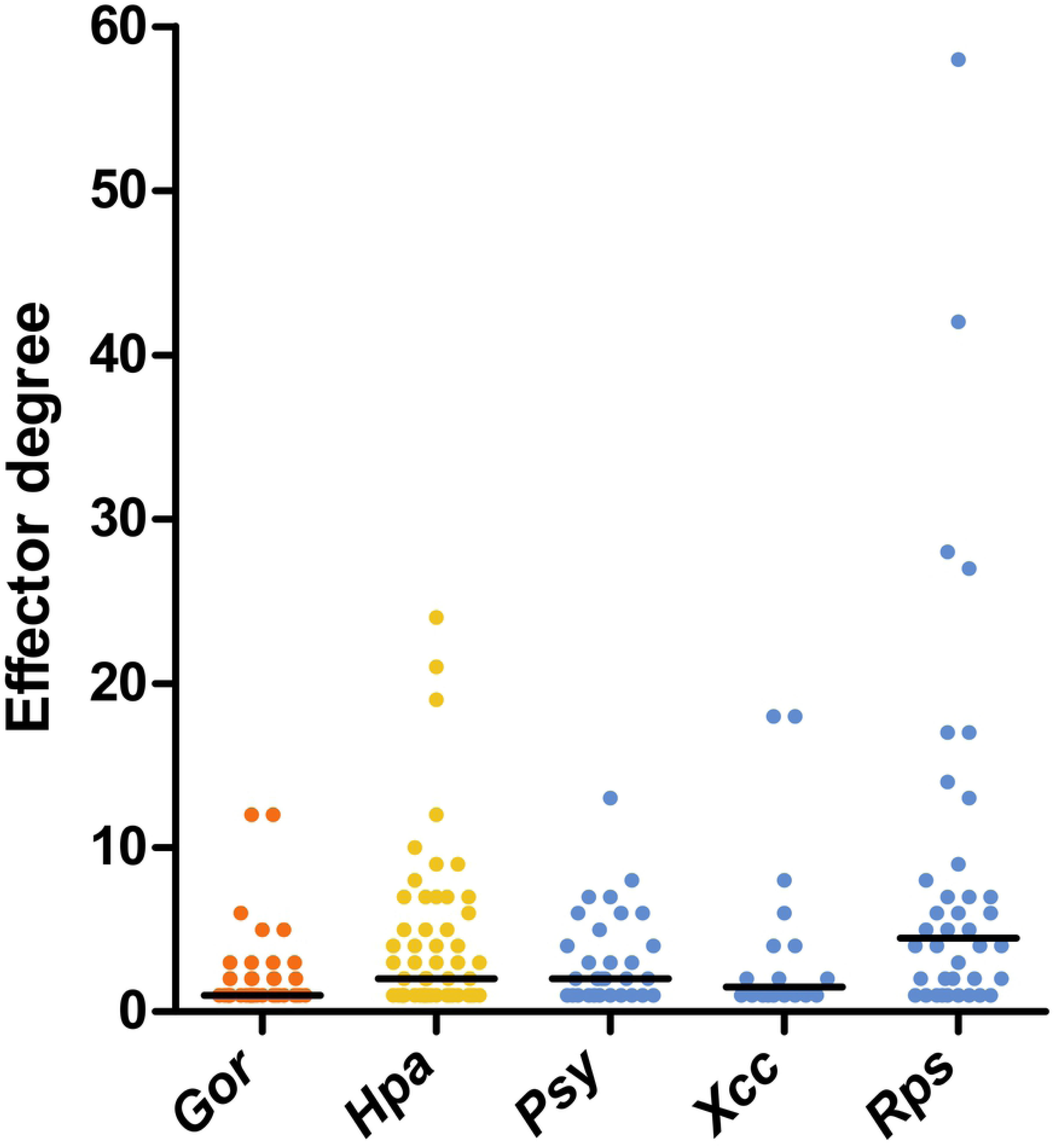
*Ath* degree of effector proteins from *Gor, Hpa*, *Psy*, *Xcc* and *Rps*. Comparison of the *Ath* degree (i.e., number of *Ath* targets per effector) of effector proteins from *Gor*, *Hpa*, *Psy*, *Xcc* and *Rps* found in the 8,000-*Ath*-cDNA collection (8K space). Horizontal black bars represent the median. Colors represent the kingdom (orange: Fungi, yellow: Chromista and blue: Bacteria).

### Effectors converge onto a limited set of Arabidopsis proteins

We compared the *Rps* and *Xcc* effector targets identified in our screenings with the targets previously identified for *Hpa*, *Psy* and *Gor* effector proteins [11, 12]. To avoid bias related to the size of the screened library, we considered only the subset of effector targets present in the 8K space (S2 Fig). At the kingdom level, the highest target specificity was found in Bacteria with 158 exclusive out of a total of 217 targets (72.8%) followed by Chromista, with 31 out of 117 (51.7%), and Fungi, with 16 out of 45 (35.6%). In total, 235 out of 299 effector targets (78.6%) were kingdom-specific. At the species level, when comparing all five pathogens, the percentage of exclusive targets was 58.9% for *Psy*, 58.7% for *Rps*, 51.7% for *Hpa*, 48.8% for *Xcc* and 35.6% for *Gor*. The total number of species-specific effector targets was 221 out of 299 (73.9%). These data show that most effector targets are kingdom- and species-specific.

To evaluate whether *Rps* and *Xcc* effectors interact randomly or converge onto a common set of *Ath* protein we performed simulations rewiring effector-*Ath* protein interactions. In these simulations, each effector was assigned randomly as many *Ath* proteins as it had interacted with in our screenings. Then, the number of targets found on all simulations was plotted and compared with the experimental data (Fig 2A). The number of effector targets observed in our screenings was significantly lower than the numbers obtained in the random simulations for both *Rps* and *Xcc*. Similar results had been reported for effectors from *Hpa*, *Psy* and *Gor* [11, 12]. This shows that, similarly to other species, both *Rps* and *Xcc* effectors also interact with a common subset of *Ath* proteins (i.e., intraspecific convergence).

**Fig 2.**
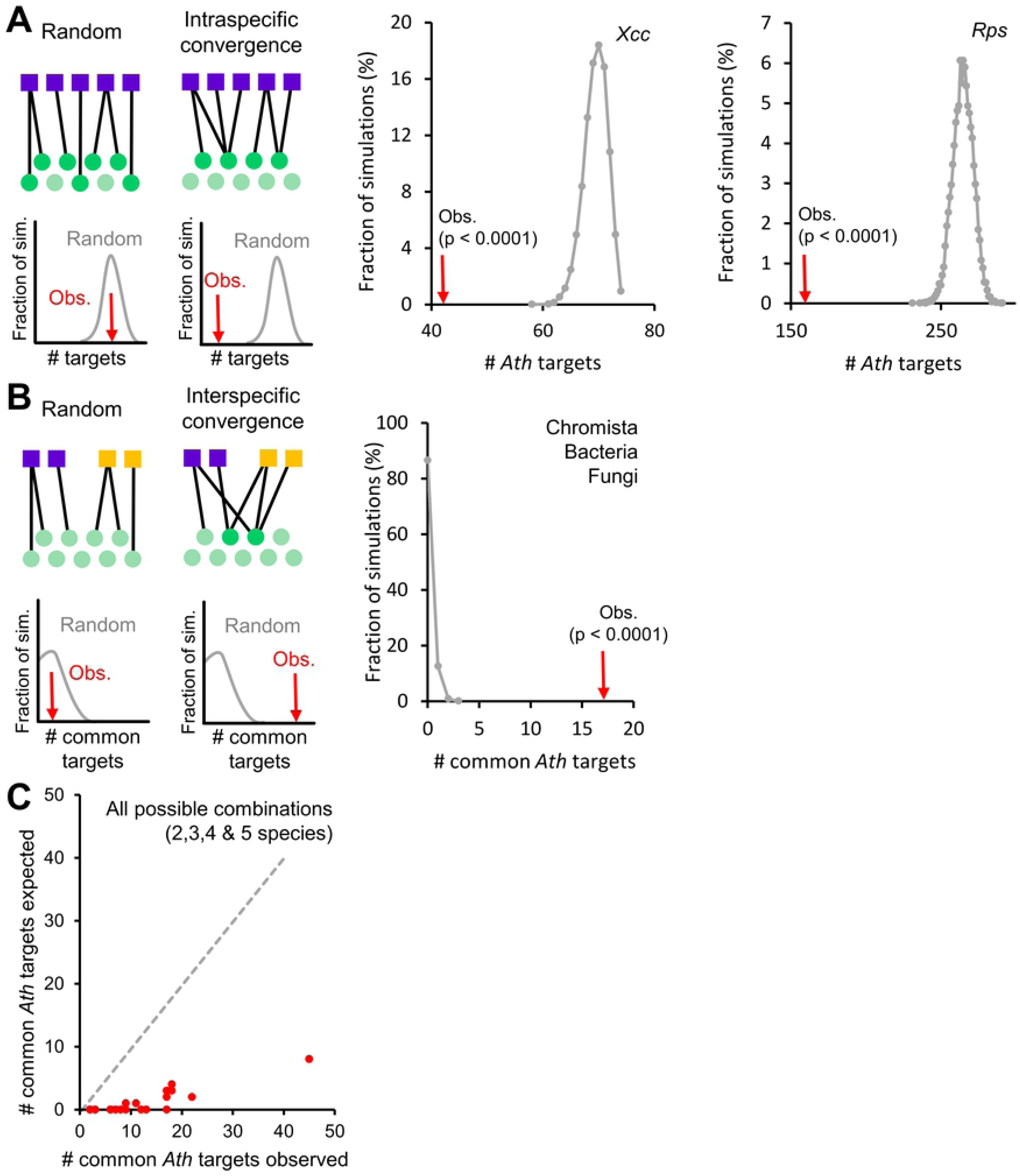
Effectors converge intra- and interspecifically onto a common set of *Ath* proteins. (A) Left: random and intraspecific convergent interactions of effectors (purple squares) with *Ath* proteins (green circles) can be distinguished by random network rewiring and simulation. Adapted from Weßling et al. [12]. Middle and right: number of *Ath* targets in the 8K space of effectors from *Xcc* strain 8004 and *Rps* strain GMI1000 found in 10,000 degree-preserving simulations (grey) versus the observed number (red arrow). (B) Left: random and interspecific convergent interactions of effectors from different species (purple and orange squares) with *Ath* proteins (green circles) can be distinguished by random network rewiring and simulation. Right: number of common *Ath* targets in the 8K space of effectors from Chromista, Bacteria and Fungi found in 10,000 simulations (grey) versus the observed number (red arrow). (C) Scatterplot of observed versus simulated number of common *Ath* targets between all binary, ternary, quaternary and quinary combinations of species. x=y regression is represented with a dashed grey line.

These random rewiring simulations also allowed us to determine whether effectors from different species interact randomly or convergently with *Ath* proteins. For this, the number of common interactors of effectors from different species was compared with the experiment data (Fig 2B). When comparing all three kingdoms, the number of common targets observed was significantly higher than expected by random rewiring. We then analyzed all possible binary, ternary, quaternary and quinary combinations of species and in all cases, the number of common targets observed was higher than expected randomly (Fig 2C). These differences were all statistically significant except for the common targets of effectors from *Psy* and *Xcc* (p-value = 0.0579) (S3 Fig). This could indicate that these two species are the most different in terms of effector targeting. However, considering that *Psy* and *Xcc* are precisely the two species with the lowest number of effectors for which targets have been identified (*Psy*: 32 and *Xcc*: 18 effector proteins), it is likely that the high p-value is caused by the limited sample size. This shows that effectors from all these five species interact with a common subset of *Ath* proteins (i.e., interspecific convergence).

Altogether, our data indicate that *Rps* and *Xcc* effectors converge both intra- and interspecifically onto a set of limited *Ath* proteins, behaving similarly to effectors from other previously screened pathogen species. This suggests the existence of a convergent set of effector targets common to evolutionary distant pathogens that might have a predominant role in the general modulation of the host responses.

### Manual curation of the literature to gather new Arabidopsis-effector protein interactions

In order to gather more knowledge on *Ath*-effector protein interactions, we conducted an extensive literature search compiling data from a wider spectrum of bacterial, fungal, oomycete and animal effector proteins. We only considered published direct protein-protein interactions that had been confirmed by classic techniques such as Y2H, co-immunoprecipitation, pull-down, protein-fragment complementation, fluorescence resonance energy transfer or mass spectrometry. We compiled 287 interactions found in 80 peer-reviewed publications involving 218 *Ath* proteins and 72 effectors from 22 pathogen species (S2 Table). Among these 22 pathogens, there were nine bacterial species, mostly proteobacteria but also a phytoplasma species; eight animal species including both nematodes and insects; four oomycete and one fungal species. While this collection of species does not represent the full diversity of *Ath* pathogens, it covers the majority of pathogens for which effector targets have been found. We can notice that despite being one of the major pathogen classes, few studies have described fungal effector interactors. This illustrates one of the current gaps in our knowledge of effector targets.

### Identification of 100 new effector hubs

To compare experimental and published data, we combined all the interactions curated from the published data together with data from our large-scale Y2H screenings. This resulted in a total of 564 different *Ath* proteins targeted by pathogen effectors. Our screenings on *Rps* and *Xcc* effectors identified 235 targets. Similar published screenings on *Psy*, *Gor* or *Hpa* effectors had identified 200 targets [11, 12]. The literature curation allowed us to identify 218 effector targets. From the 235 *Rps* and *Xcc* effectors targets found in our screening, 166 were new which represents 29.4% of the total targets compiled in this study (Fig 3). This highlights the potential of such systematic and high throughput large-scale screenings in identifying novel effector targets. The average effector degree (i.e., number of effectors interacting with an *Ath* protein) was 2.3 but it was unevenly distributed among the 564 targets with 350 of them interacting with only one effector (62%) and 14 interacting with more than 10 effectors (2.5%) (S4 Fig). The contribution of our experimental data was important in the identification of single targets as we identified 93 out of the 350 (26.6%). More remarkable was our contribution in the identification of “effector hubs”, what we defined as *Ath* proteins interacting with two or more effectors (Fig 4). The definition of hub has been debated and it has been traditionally associated with proteins that are highly connected in interactomic networks [19]. Our definition of “effector hub” came from the need to designate the *Ath* proteins that interact with several effectors and is based exclusively on the number of interacting effector proteins. We identified 100 new effector hubs and increased the degree of 42 previously described effector hubs (S3 Table).

**Fig 3.**
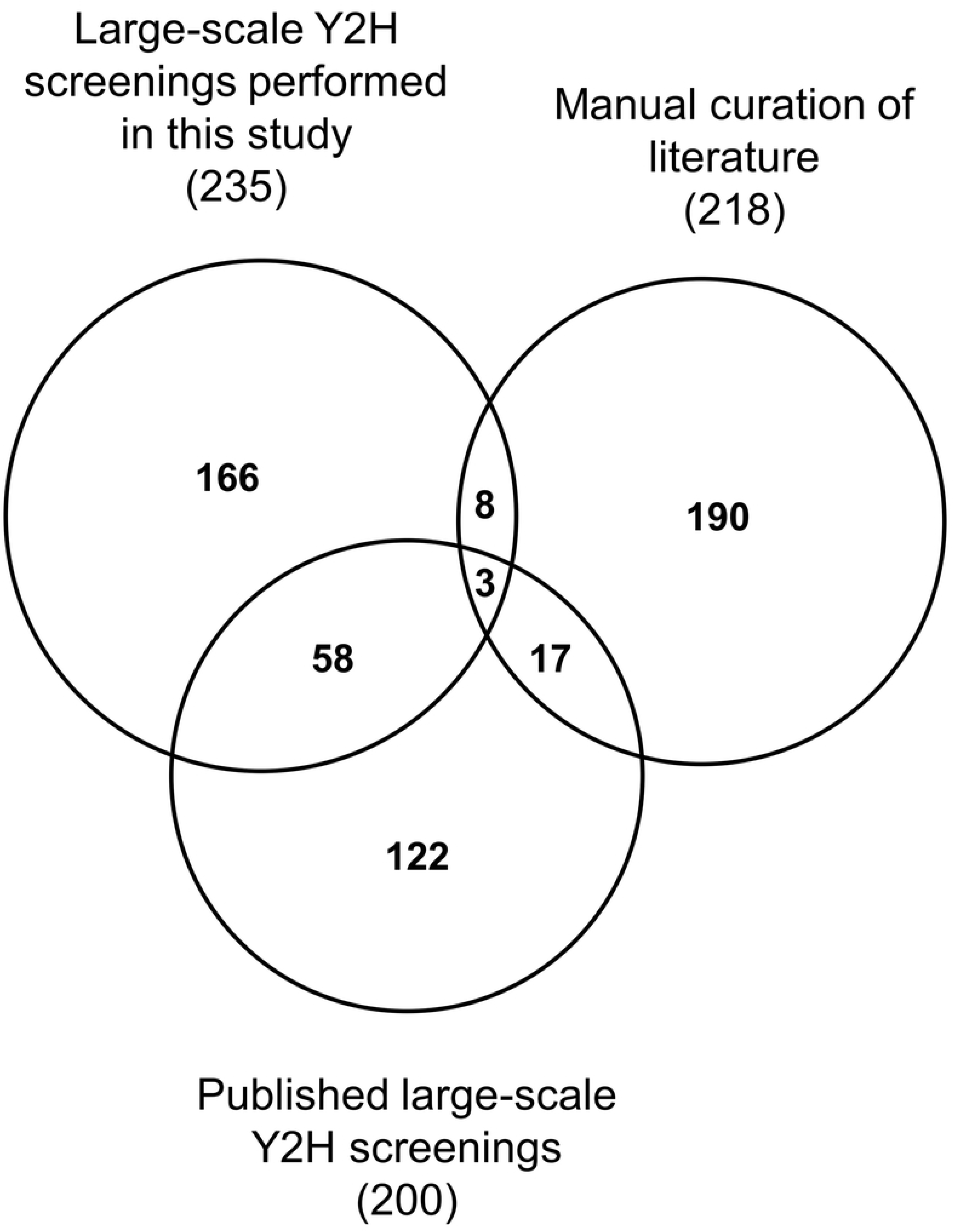
Overlap among effector targets depending on the origin of the dataset. Area-proportional Venn diagram showing the overlap among effector targets identified in the large-scale Y2H screenings performed in this study, in similar large-scale Y2H already published [11, 12] and in the manual curation of the literature. The total number of effector targets coming from each dataset is indicated in brackets.

**Fig 4.**
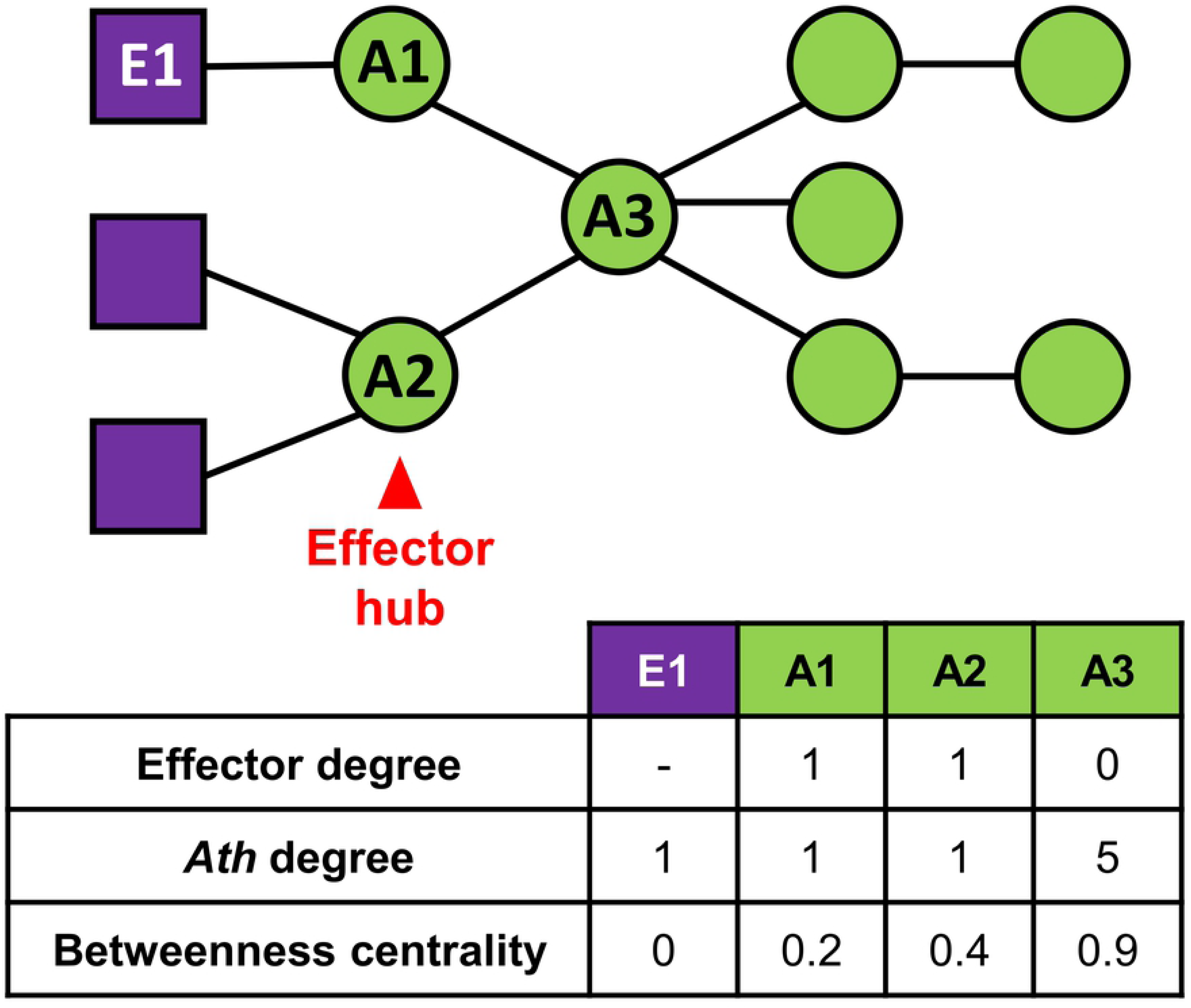
Network topology parameters. Example of a simple interactomic network of three effector proteins (purple squares) and eight *Ath* proteins (green circles) to illustrate our definition of “effector hub” (i.e., *Ath* protein interacting with two or more effectors; highlighted in red) and the three network topology parameters analyzed in this study. 1) Effector degree: number of effectors that interact with a given *Ath* protein. 2) *Ath* degree: number of *Ath* proteins that interact with a given effector or *Ath* protein. 3) Betweenness centrality: fraction of all shortest paths connecting two proteins from the network that pass through a given protein.

To evaluate the potential relevance of the newly identified effector hubs in plant immunity, we conducted a second literature survey to check if the corresponding *Ath* genes had been previously characterized to be involved in plant immunity or pathogen fitness *in planta* (Table 1). 16 out of the 100 new effector hub genes, have already been described for their altered infection or other immunity-related phenotype when mutated, silenced or overexpressed. Additionally, the orthologs of 3 other new hubs in other plant species, also produced altered infection phenotypes when silenced or overexpressed. A total of 19 out of the 100 newly identified effector hubs have already been shown to be involved in biotic stress responses. Considering that many of the remaining newly defined effector hubs have been poorly characterized (e.g., hypothetical proteins or descriptions based on homology or belonging to a protein family), it is likely that the number of effector hubs involved in immunity was underestimated. This constitutes a valuable source of novel candidates for further functional characterization.

**Table 1.**
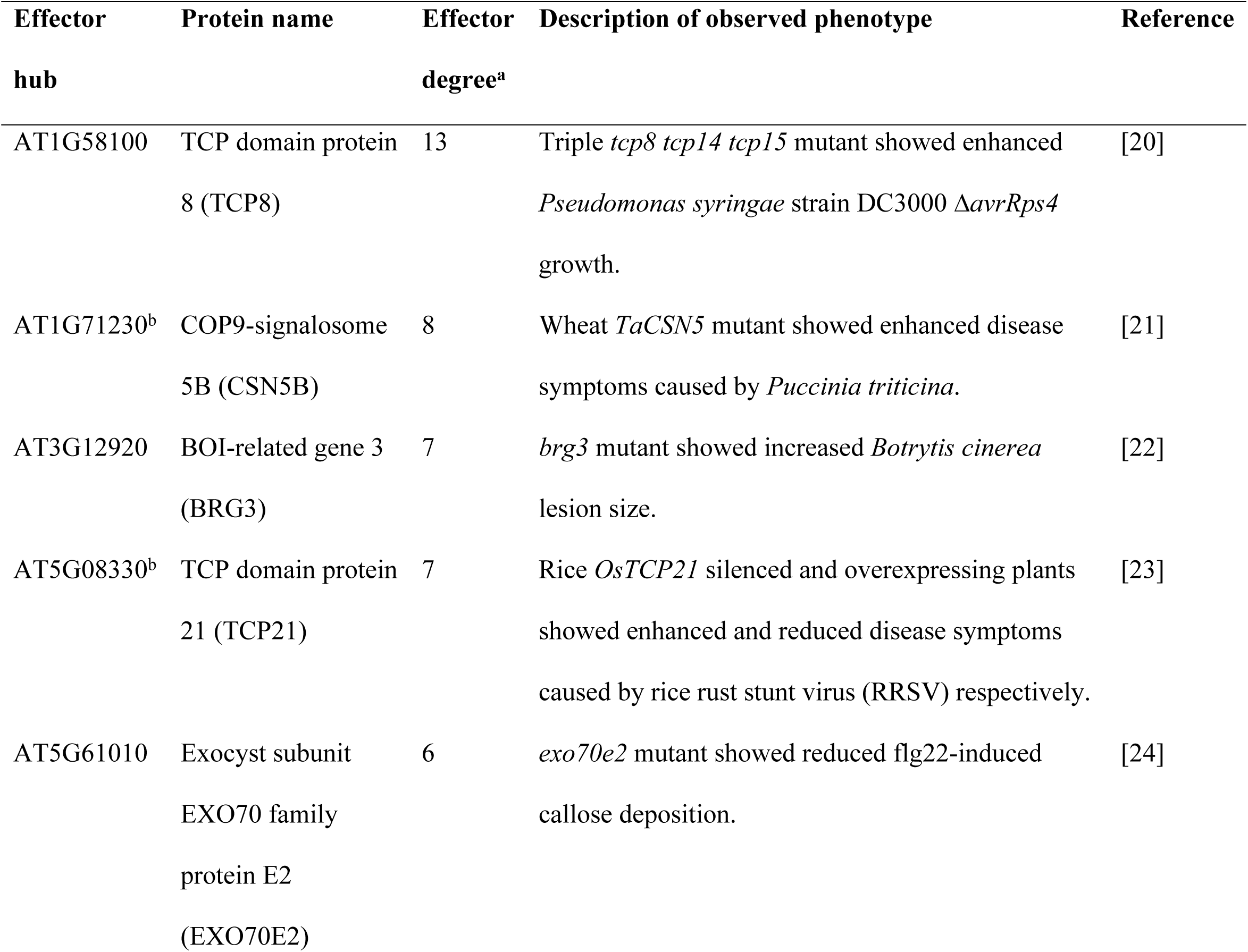
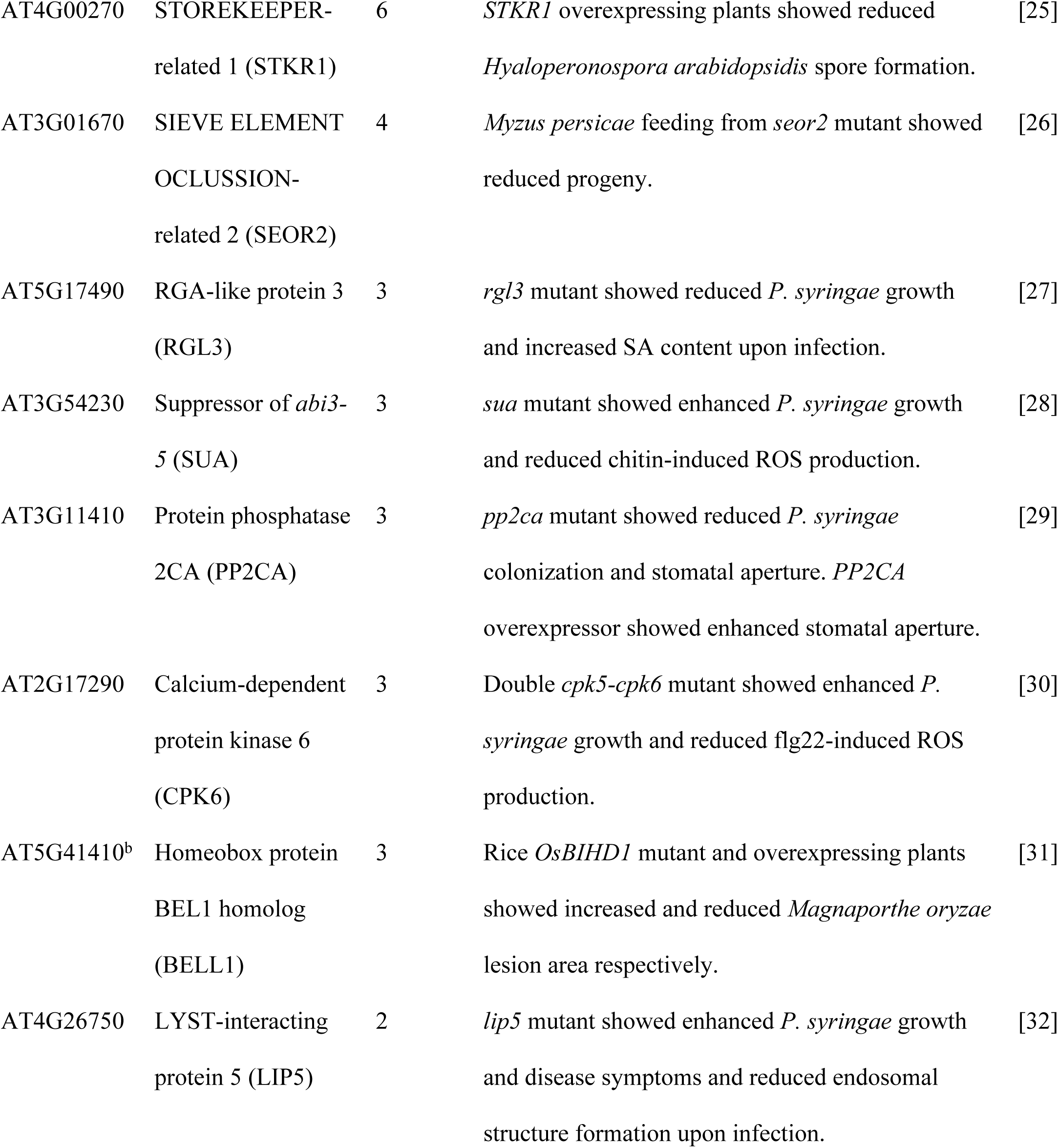
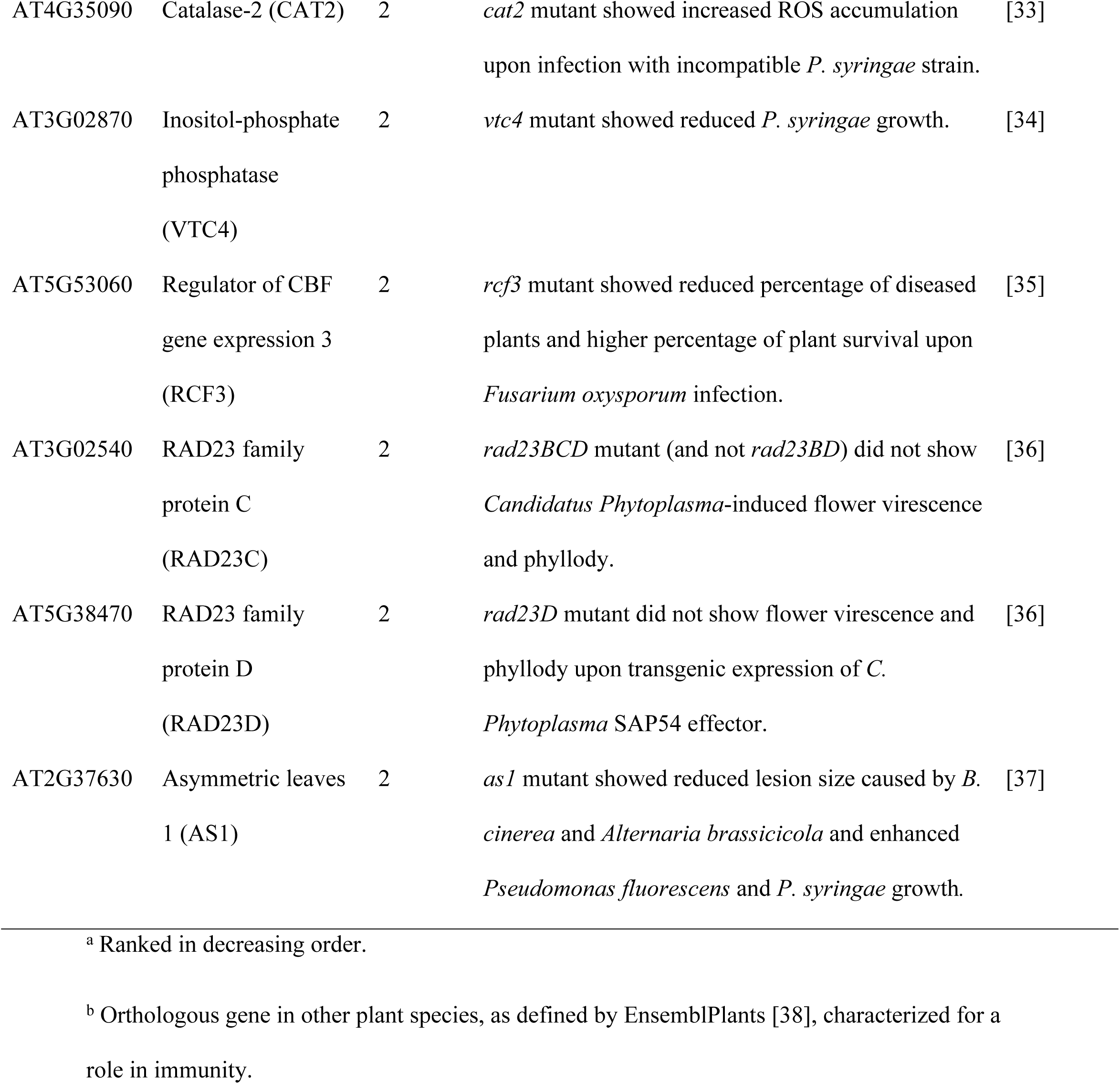
List of 19 new effector hubs involved in plant immunity.

In terms of organism of origin, most of the 564 targets are bacterial effector targets as it could be expected considering that 132 out of the 266 total effectors compiled came from bacteria (S4 Fig). In the case of effector hubs, it is noteworthy that 133 out of the 214 hubs described in this work are targeted by effectors from a single kingdom while there are only 64, 16 and 1 hubs interacting with effectors from 2, 3 or 4 different kingdoms respectively. Although biased by the structure of the data, this could suggest kingdom specificity of effector targeting.

### Effector targets tend to occupy key positions for the network organization

We constructed an *Ath*-effector protein interaction network compiling the previously described experimental and literature-compiled data with *Ath*-*Ath* protein interactions from public databases and the literature [39–42]. From the total of 49,500 interactions compiled in this study, 48,597 were grouped into a single connected component constituting what we defined as our *Ath*-effector interactomic network (Table S4). This network was constituted of 47,314 *Ath*-*Ath* and 1,283 *Ath*-effector protein interactions between 8,036 *Ath* proteins and 245 effector proteins. Effectors came from 23 different species including bacteria (128 effectors), oomycetes (61 effectors), fungi (46 effectors) and animals (10 effectors). The uneven distribution of effectors among kingdoms highlights the contribution of the large-scale screenings in the identification of effector targets as 1,002 out of 1,283 *Ath*-effector protein interactions came from either our experimental data or previous screenings of the same library [11, 12].

To further investigate the potential impact of effectors on the plant interactome, we evaluated the importance of their targets for the organization of the network. We focused on two main network topology parameters: degree and betweenness centrality (Fig 4). The degree of a protein represents the number of proteins that it interacts with. In this study we differentiated two types of degrees depending on the nature of the interacting proteins: the *Ath* degree of a given effector or *Ath* protein (i.e., number of interacting *Ath* proteins) and effector degree for a given *Ath* protein (i.e., number of interacting effector proteins). The betweenness centrality of a protein is the fraction of all shortest paths connecting two proteins from the network that pass through it. There are two main types of key proteins in a network [43]: 1) proteins important for local network organization, typically showing high degree, and 2) proteins important for the global diffusion of the information through the network, characterized by high betweenness centrality. It had been previously reported in more limited networks that effectors tend to target host proteins with high degree and centrality [43–45]. We then analyzed whether this was the case in our network comparing effector targets with the rest of the *Ath* proteins (Fig 5). The fraction of proteins decreased rapidly as the Ath degree increased. This indicates that most *Ath* proteins present low *Ath* degree and only a few of them show high *Ath* degree values. This tendency was significantly shifted towards higher *Ath* degree values in effector targets compared to the rest of *Ath* proteins. To represent this tendency shift we estimated and compared the area under the curve values of the cumulative distribution of *Ath* degree for effector targets and the rest of *Ath* proteins (Table 2). Effectively, the area under the curve value of effector targets was higher than the value of the rest of *Ath* proteins. This indicates that effector targets present generally higher *Ath* degree than the rest of *Ath* proteins. Similarly, we compared the betweenness centrality of these two groups of proteins (Table 2 and Fig S5). Effector targets also presented significantly higher betweenness centrality values than the rest of *Ath* proteins. Altogether, these results indicate that effectors preferentially interact with *Ath* proteins that are more connected to other *Ath* proteins and that occupy more central positions in the interactomic network.

**Fig 5.**
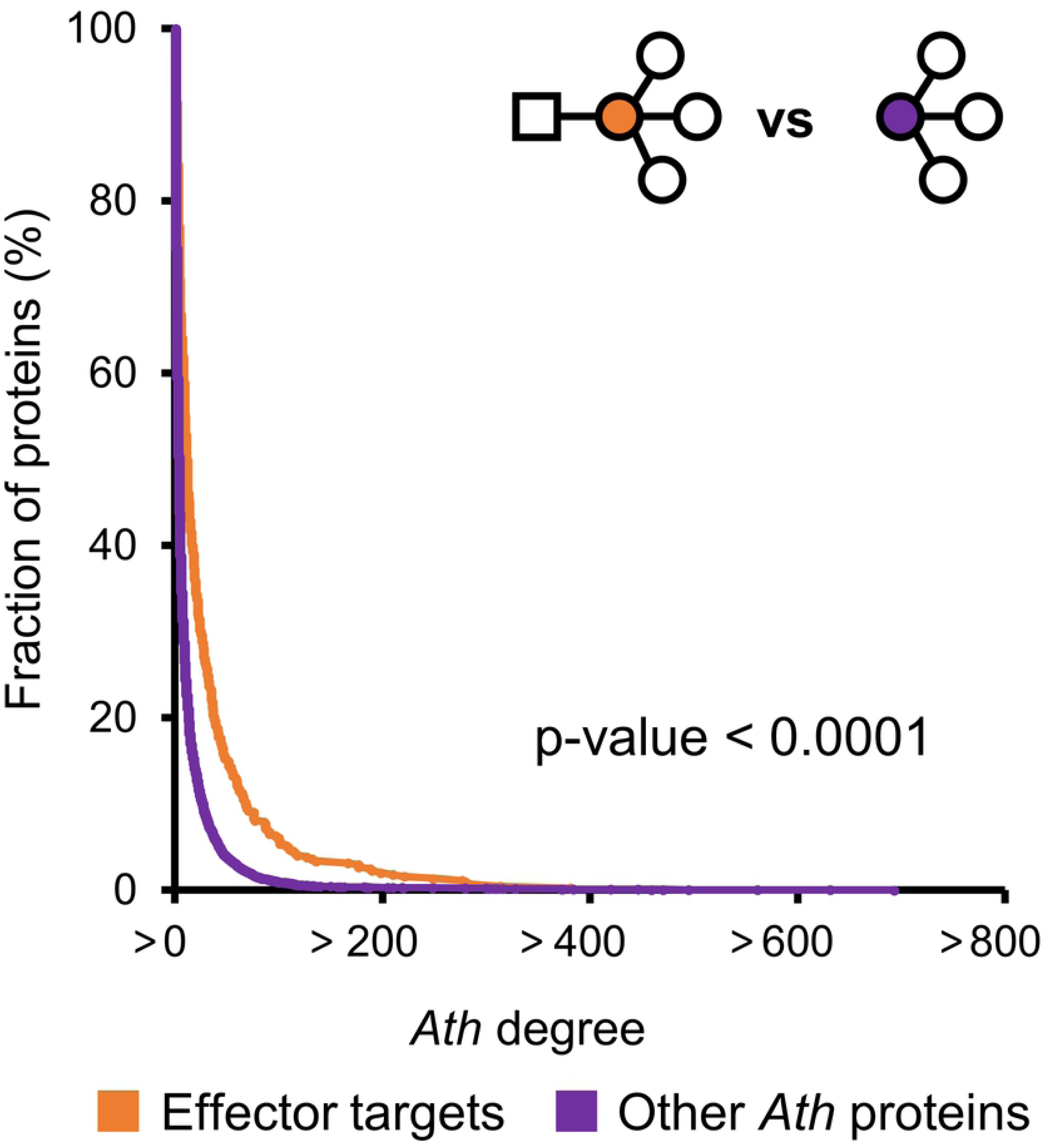
*Ath* degree of Ath proteins targeted or not by effectors. Cumulative distribution of *Ath* degree of *Ath* proteins targeted (orange) or not (purple) by effectors. The significance of the difference was validated by one-tailed Wilcoxon signed-rank test. The illustration in the upper right corner represents each compared group. Effectors are represented by squares, *Ath* proteins by circles and the color code matches the cumulative distribution graph.

**Table 2.**
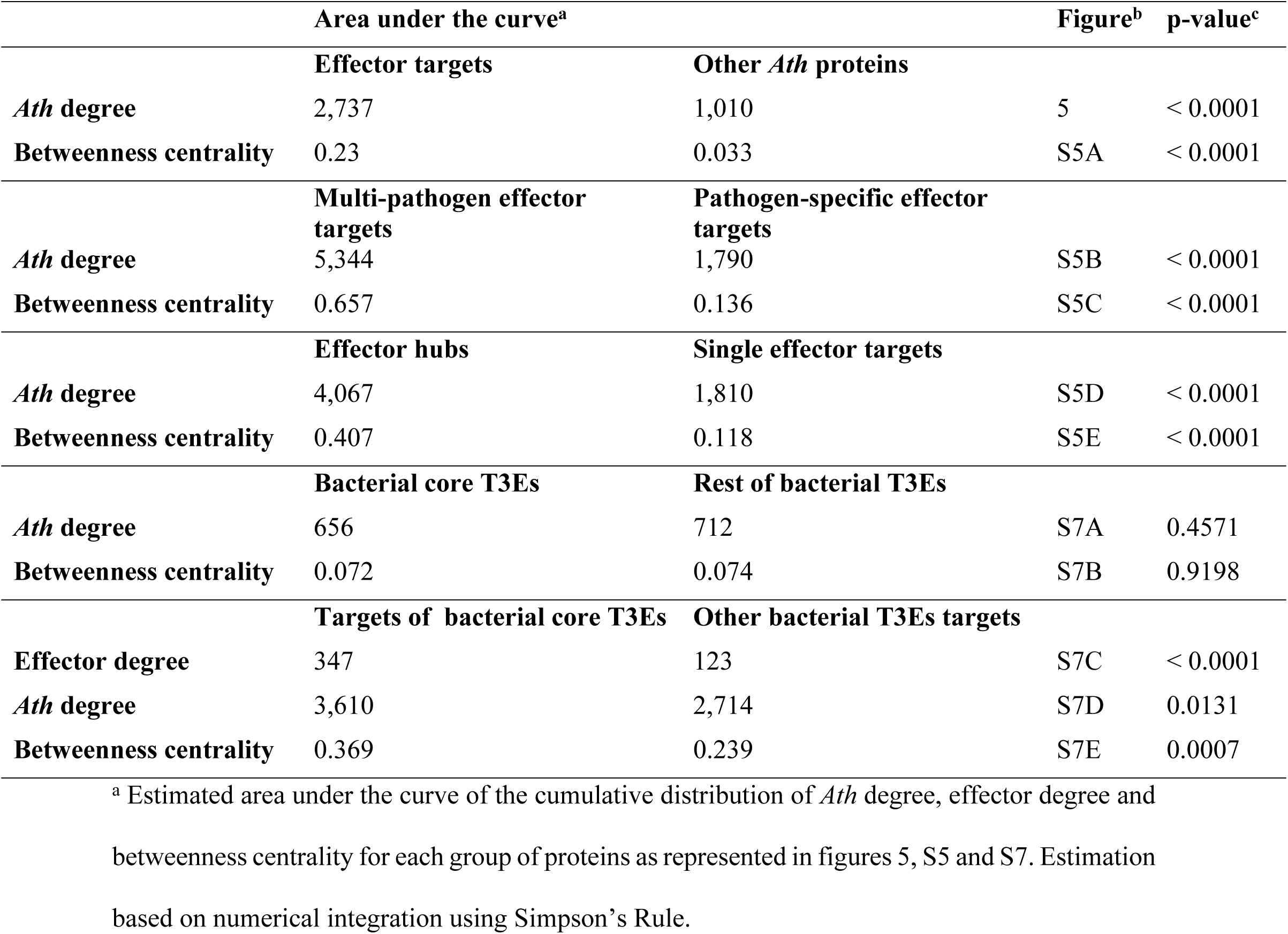
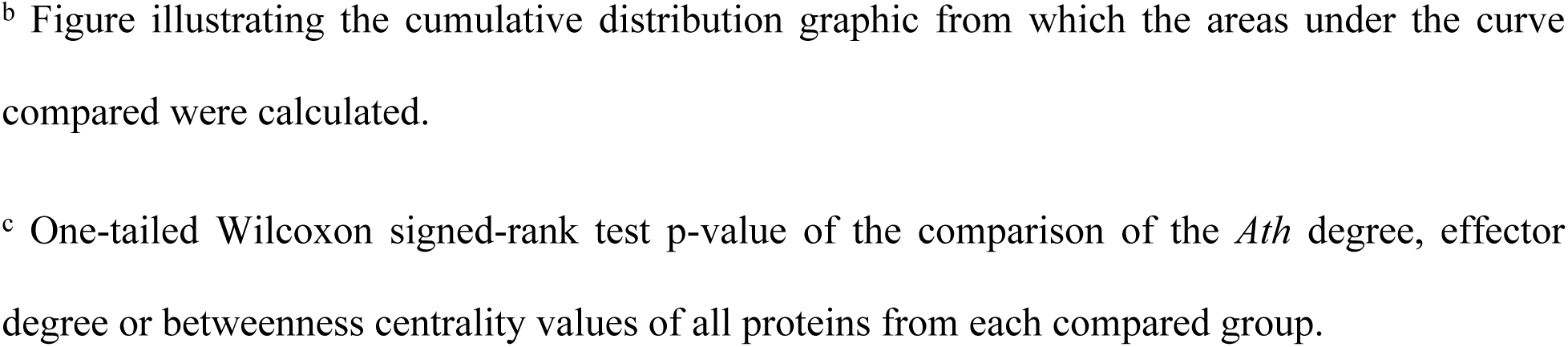
Cumulative *Ath* and effector degrees and betweenness centrality of different groups of effector targets

We then wanted to test if the *Ath* degree and betweenness centrality values differed among distinct types of effector targets (Table 2 and Fig S5). First we compared multi-pathogen and pathogen-specific targets as previously described (S2 Fig). Multi-pathogen effector targets presented significantly higher *Ath* degree and betweenness centrality compared to pathogen-specific effector targets. We also compared effector hubs with single effector targets. Similarly, effector hubs also showed higher betweenness centrality and *Ath* degree than single effector interactors. This last observation implies that an *Ath* protein that interacts with several effectors tends to interact with more *Ath* proteins as well. To evaluate whether this is biologically relevant or a bias of the ‘stickiness’ of a protein, we compared the *Ath* and effector degree values of all targets. Our results showed that these two parameters are not correlated (Pearson correlation coefficient = 0.3221) (S6 Fig). This suggests that effector hubs interact with more *Ath* proteins than single effector targets and not because they might be stickier.

In this work, by compiling our experimental interactomic data on *Xcc* and *Rps* and the literature-curated interactions from a wide variety of other pathogen effectors, we extended the notion that effectors tend to privilege interactions with host proteins with higher *Ath* degree and betweenness centrality [43, 45]. Furthermore, we showed that this tendency is stronger among effector hubs compared to single targets and among multi-pathogen effector targets compared to pathogen-specific targets. This reflects the importance of interfering with these key position proteins in the modulation of host-pathogen interactions.

### Bacterial core T3Es interact with more connected and central *Ath* proteins

Our work on *Rps* and *Xcc* together with previous work on *Psy* T3Es [11] provided a large amount of interactomic data on bacterial pathogen species for which other resources have been generated, particularly in terms of abundance and diversity of sequenced genomes and thus curated T3E repertoires [18,46–50]. The most conserved set of T3Es, or ‘core effectome’, from each of the three bacterial species has been previously defined [47,48,50]. We then tested whether these subsets of T3Es behaved differently from the rest of bacterial T3Es in terms of interaction with host proteins (Table 2 and Fig S7). Our data showed that core and variable T3Es from the three species do not differ in *Ath* degree nor betweenness centrality. We then tested if there were any differences between the network properties of the targets of core T3Es and the other bacterial T3E targets. Core T3Es targets showed higher effector degree, *Ath* degree and betweenness centrality than the rest of targets of bacterial T3Es. This suggests that, although core T3Es in general do not have more targets than the rest of bacterial T3Es, they do interact with more highly connected and central *Ath* proteins. This might imply that core T3Es have a larger potential to interfere with the host interactome what could explain the selective pressure to maintain them in the majority of strains.

### EffectorK, an online interactive knowledge database to explore the Arabidopsis-effector interactomic data

In order to facilitate the access and exploration of all the data presented in this work, we have generated EffectorK (for ‘Effector Knowledge’), an interactive web-based knowledge database freely available at www.effectork.org. The latest version (October 2, 2019) contains 49,875 interactions 8,617 proteins coming from 2,035 publications. From these, 1,300 are *Ath*-effector protein interactions. Searches can be done based on a wide range of supported identifiers such as different protein names, NCBI or TAIR accession numbers, PubMed identifiers and InterPro terms. Additionally, users can also query nucleotide or amino acid sequences directly with BLAST or use accession numbers from other model and crop plants to find homologs within the database. All proteins found by query are then listed in tabular format and hyperlinked to the corresponding interactomic data, external resources and amino acid sequences. Interactomic data for a given protein can be then explored and downloaded in graphical or tabular format. The visual interface for the graphical representation of the interactomic data allows users to expand or re-center a local subnetwork based on a given protein, get information and access to external resources linked to either a protein (node) or an interaction (edge) or modify the layout and the position of the elements for optimal visualization. Additionally, EffectorK also allows users to find the shortest paths between two queried proteins in the network.

In order to update, expand and further improve EffectorK, we encourage users to submit their own interactomic data by filing in and sending the dedicated template. These data will be verified by the curator team prior to their incorporation in the database. More information about usage, content and data submission is accessible online, under the tabs ‘Help’ and ‘Contribute’ of the database web server. Please contact us if you have any question or suggestions by email at.

## Discussion

In this study we identified systematically *Ath* targets of effectors from the vascular bacterial pathogens *Rps* and *Xcc*. We combined this information with other *Ath* targets identified in similar experimental setups. Additionally, we conducted an extensive literature review to gather published *Ath* targets of effectors from a wide variety of pathogens including other bacterial species and also oomycete, fungal and animal pathogens. Studying this combined interactomic dataset allowed us to identify new trends of how effectors interfere with the plant proteome and evaluate whether previously described network principles were still supported on a wider scale. We showed that there are no substantial differences in terms of connectivity among the effectomes of five different pathogen species screened systematically (Fig 1). We have reinforced previously described intra- and interspecific convergence of effector targeting with effectors from two new species [11, 12], and showed at the same time that most effector targets are pathogen specific (Fig 2 and S2). Our analyses also supported the previously described tendency of effectors to interact with plant proteins better connected and central in the network [43, 45], and showed that this tendency is even stronger among effector hubs, multi-pathogen targets and bacterial core T3E targets (Table 2 and Fig S5).

### The balance between target specificity and convergence

Our data showed that most effector targets were pathogen-specific (S2 Fig) but at the same time, effectors converge interspecifically onto a small subset of *Ath* proteins (Fig 2B-C). These *a priori* contradictory observations open an interesting question: what is the balance between specificity and convergence of effector targets? At this point, it is impossible to assert whether this specificity is merely caused by the limited number of pathogens screened at the effectome-scale or if it is a reflection of the different and unique ways that each pathogen has evolved to interfere with the host physiology and immunity. This issue can only be addressed by increasing the number of pathogen effectors screened thoroughly and at a large-scale. Comparing large datasets of effector targets of a wider and more diverse set of pathogens would allow evaluating in which sense this balance between specificity and convergence tilts: 1) If the target specificity decreased, it would mean that the effectomes from the different pathogens tend to interact similarly with the host proteome. This was the case when we compared the percentage of species-specific targets of effectors from *Hpa*, *Psy* and *Gor* that passed from being 73.9%, 64.9% and 46.7% in previous works [11, 12], to 51.7%, 58.9% and 35.6%, respectively in the present study (S2 Fig). Nevertheless, a total of five screened species is probably not powerful enough to sustain this claim. 2) If, on the contrary, the target specificity increased with the number of screened species, it would mean that the different pathogens have evolved unique ways to modulate the interaction with the host. If this were be the case, deeper analyses comparing related pathogens (e.g., species with similar lifestyle or from the same kingdom) could allow identifying trait-specific targets (e.g., effector targets exclusive among vascular pathogen effectors). In any case, to better understand the similarities and particularities on how effectors modulate host processes, it is essential to increase the number of pathogen species screened for effector targets at the effectome-scale.

### Large-scale screenings fill the gap in the identification of effector targets

Including manually curated data from literature has allowed us to broaden significantly the diversity of plant pathogen species compared to similar studies. However, 346 out the 564 described *Arabidopsis* effector targets have been identified exclusively through large-scale Y2H screenings against partial libraries of *Ath* cDNAs. As with any other large-scale screening, the technical limitations together with the incompleteness of the library might have probably led to an underestimation of the plant-effector interactome of the five screened species [51]. The relatively small overlap between the large-scale Y2H screenings and manually curated literature datasets might be a consequence of this limitation (Fig 3). This small overlap illustrates the current knowledge gap in the characterization of the full plant interactome of pathogen effectors. Extensive work will be required to characterize further effector-host protein interactions in other pathosystems. As one of the simplest yet powerful high throughput techniques for protein-protein interaction detection, our work, like others before, highlights the potential of such large-scale Y2H screenings in the identification of novel effector targets in an easy, cheap and systematic manner.

### EffectorK, an entry point to explore and make sense of plant-effector interactomics

To conclude, our work also provides valuable resources for the plant-pathogen interaction community. We described 540 new *Ath*-*Rps* and *Ath-Xcc* effector protein interactions that allowed us to identify 166 new effector targets (S1 Table). We also manually curated several publications to assemble a collection of 287 *Ath*-effector protein interactions from a wide variety of pathogens (S2 Table). All this, allowed us to identify 100 novel effector hubs (S3 Table). The contribution to plant immunity of these effector hubs has been described for 19 of them, but remains untested for the majority (Table 1). This constitutes a list of promising candidates for further functional characterization. All these data were integrated in EffectorK, a knowledge database where users can have easy access to the *Ath*-effector protein interactions and explore the resulting interactomic network visually and interactively. While major efforts were done to capture the maximal diversity on the pathogen side, we limited our work to the Arabidopsis plant model. Thanks to the built-in homology search tools available, users can also use their own data as query regardless of the species studied. It is therefore feasible to use EffectorK as a starting point to build on and extend to crop plant-effector protein interactomics. In the long term, these data could be exploited to better understand how pathogens interact with these crops with the prospect of selecting breeding candidates for improved tolerance or resistance against pathogens.

## Materials and Methods

### Cloning of *Rps* and *Xcc* T3E genes

All the cloning of the T3E genes from *Rps* and *Xcc* was performed by BP gateway BP or TOPO cloning (Thermo Fisher Scientific), to generate pENTRY plasmids, which were later transferred into the appropriate Y2H plasmids [11], using the LR gateway reaction (Thermo Fisher Scientific). S5 Table contains all the PCR primers and final plasmid identities describing the collection of plasmids used in this study. Gene sequence information from *Rps* strain GMI1000 can be obtained from www.ralsto-T3E.org [47] and from the published genome of *Xcc* strain 8004 [52].

### Y2H screenings

The Y2H screening was performed in semi-liquid (‘8K space’ screening) and liquid (‘12K space’ screening) media as recently reported [53], which is an adaptation of a previously developed Y2H-solid pipeline [54]. In both protocols the same low copy number yeast expression vectors and the two yeast strains, *Saccharomyces cerevisiae* Y8930 and Y8800, were used. The expression of the *GAL1-HIS3* reporter gene was tested with 1 mM 3AT (3-amino-1,2,4-triazole, a competitive inhibitor of the HIS3 gene product), unless described otherwise. Prior to Y2H screening, DB-X strains were tested for auto-activation of the *GAL1-HIS3* reporter gene in the absence of AD-Y plasmid. In case of auto-activation, DB-X were physically removed from the collection of baits and screened against the (DB)-*Ath*-cDNA collections using their AD-X constructs. Briefly, DB-X baits expressing yeasts were individually grown (30°C for 72 hours) into 50-ml polypropylene conical tubes containing 5 ml of fresh selective media (Sc-Leucine, Sc-Leu). Pools were created by mixing a maximum of 72 and 50 individual bait yeast strains for the ‘8K space’ and ‘12K space’ respectively. Subsequently, 120 µl and 50 µl of these individual pools were plated into 96-well and 384-well low profile microplates for *Ath*-cDNA ‘8K space’ and ‘12K space’ collections respectively. Glycerol stocks of the (AD)-*Ath*-cDNA ‘8K space’ and ‘12K space’ collections were thawed, replicated by handpicking or using a colony picker Qpix2 XT into 96-well and 384-well plates filled with 120 µl and 50 µl of fresh selective media (Sc-Tryptophan; Sc-Trp) respectively, and incubated at 30°C for 72 hours. Culture plates corresponding to the DB-baits pools and AD-collection were replicated into mating plates filled with YEPD media and incubated at 30°C for 24 hours. In liquid Y2H case (‘12K space’ screening), mating plates were then replicated into screening plates filled with 50 µl of fresh Sc-Leu-Trp-Histidine + 1 mM 3AT media and incubated at 30°C for 5 days. In order to identify primary positives, the OD_600_ of the 384-well screening plates was measured using a microplate-reader Tecan Infinite M200 PRO. In semi-liquid Y2H case (‘8K space’ screening), mated yeast were spotted onto Sc-Leu-Trp-Histidine + 1 mM 3AT media agar plates, and incubated at 30 °C for 3 days. Protein pairs were identified by depooling of DB-baits in a similar targeted matricial liquid or semi-liquid assays in which all the DB-baits were individually tested against all the previously identified AD-proteins. Identified pairs were picked and checked by PCR and DNA sequencing.

### Database content and manual curation

Binary interactions between *Ath* proteins with each other and with pathogen effector proteins were compiled on tabular form keeping track of the protein names and accessions, species and ecotypes/strains of origin, techniques used to detect the interactions and the reference. *Ath*-*Ath* protein interactions were compiled from the Arabidopsis Interactome [41, 42] and the public databases BioGrid (www.thebiogrid.org [39]; downloaded in September 2019) and IntAct (www.ebi.ac.uk/intact [40]; downloaded in September 2019). We only kept the direct interactions with the evidence codes ‘co-crystal structure’, ‘FRET’ (fluorescence resonance energy transfer), ‘PCA’ (protein-fragment complementation assay), ‘reconstituted complex’ or ‘two-hybrid’ on BioGrid and ‘physical association’ on IntAct. *At*-effector protein interactions were gathered from our experimental Y2H data together with the similarly produced data on *Hpa*, *Psy* and *Gor* effectors [11, 12]. In addition, an extensive keyword search on effector-Arabidopsis literature was done to retrieve interactions from 80 published articles. A confidence level was assigned to each interaction depending on the number of independent techniques used in a publication for validation: “1” if the interaction was detected by only one technique and “2” if the interaction was validated by at least a second technique. Some interactions lacked important information but, in order to maximize the extent of our network, several assumptions were taken instead of discarding useful data. First, gene models for *Ath* proteins were rarely mentioned on publications so we assumed the first gene model available on the latest version of the Arabidopsis genome (Araport11 [55]). Secondly, when the ecotype/strain of the organism was not explicitly stated, a generic ‘NA’ (not available) was assigned.

### In silico analyses

#### Computational simulations of random targeting of *Ath* proteins by single pathogen effectors (intraspecific convergence)

Significance of the intraspecific convergence was tested comparing our experimental data with random simulations as previously published [12]. Briefly, for each effector of *Xcc* and *Rps* we assigned randomly the same number of *Ath* targets as experimentally observed from the degree-preserved list of 8K proteins. The distribution obtained from 10,000 simulations was plotted and compared to the experimentally obtained data. The p-value of the experimental data was calculated as follows: number of simulations where the number of targets is lower or equal than experimentally observed is divided by the number of simulations. When the number of simulations with less targets than observed was zero, the p-value was set to < 0.001.

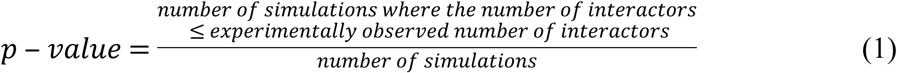

#### Computational simulations of random targeting of *Ath* proteins by several pathogen effectors (interspecific convergence)

Significance of the interspecific convergence was tested comparing our experimental data and previously published data with random simulations as published [11, 12]. Briefly, for each effector of all compared pathogens we assigned the same number of *Ath* targets as experimentally observed/published from the list of 8K proteins. The distribution obtained from 10,000 simulations was plotted and compared to experimentally and published data. The p-value of the experimental data was calculated as follows: number of simulations where the number of common targets between species was higher or equal than the experimentally observed is divided by the number of simulations. When the number of simulations with more common targets than observed was zero, the p-value was set to < 0.001.

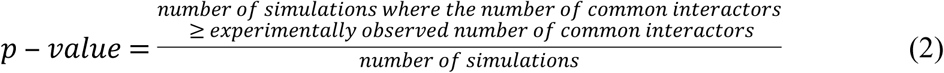

#### Overlap of effector targets

The overlaps of effector targets between the different kingdoms and species were calculated taking into account the targets found in the different large-scale screening and limiting to the 8K space. For representation of the data, Venn diagrams were generated using the Venn Diagrams tool from VIB-UGent Center for Plant Systems Biology (www.bioinformatics.psb.ugent.be/webtools/Venn/). The overlap of effector targets from the different datasets was calculated not limiting to any limited space. For an area-proportional representation of the data, a Venn diagram was generated using BioVenn [56].

#### Network topology analyses

The topology parameters of the *Ath*-effector interactomic network were calculated on Cytoscape 3.7.2 [57]. Our analyses focused on two key node parameters: degree and betweenness centrality. The degree of a protein is a measure of its connectivity and denotes the number of proteins interacting with it. Throughout this work, we have differentiated two kinds of degrees: 1) effector degree (i.e., number of interacting effector proteins) and 2) *Ath* degree (i.e., number of interacting *Ath* proteins). The betweenness centrality measures the proportion of shortest pathways between two proteins that passes through a given node. These parameters were compared against different subset of data and statistical tests were performed in R language [58]. The cumulative distribution of these parameters among different subset of data was plotted and the area under the curve was estimated using Simpson’s rule with the ‘Bolstad2’ package [59].

### Database construction

The database was built using the software architecture recently described [60]. The files submitted by the curator team were automatically checked for typographic mistakes using *ad-hoc* Perl scripts and loaded into a Neo4J database and indexed in an ElasticSearch search engine. Each release was rebuilt from scratch. Data were made accessible through a web interface (see Results and discussion section) built upon Cytoscape.js library [61]. The raw data used for the database setup are available in the ‘Data’ section of www.effectork.org and the source code is available at https://framagit.org/LIPM-BIOINFO/KGBB.

## Acknowledgments

We wish to thank Alberto Macho and Laurent Deslandes for the contribution of RipS3, and RipP1 and RipP2 containing plasmids. We thank colleagues (Corinne Audran, Julien Luneau, Effectome and FNX network members) for their critical opinion on our work.

## Supporting information

**S1 Fig. *Ath* degree of T3E proteins from *Rps* strain GMI1000 and *Xcc* strain 8004.**

*Ath* degree (i.e., number of *Ath* targets per effector) in the in the 12,000 (12K space, light blue) and 8,000 *Ath* cDNA collections (8K space, dark blue) of T3E proteins from *Rps* strain GMI1000 (A) and *Xcc* strain 8004 (B). For *Rps* strain GMI1000: in the first screening RipA3, RipAA, RipAB, RipAC, RipAG, RipAL, RipAM, RipAN, RipAO, RipAP, RipAQ, RipAR, RipAZ1, RipB, RipBA, RipG3, RipG4, RipG6, RipG7, RipH2, RipH3, RipI, RipK, RipM, RipN, RipO1, RipP1, RipQ, RipR, RipS2, RipS6, RipT, RipTPS, RipX and RipZ were screened but no targets were found. In the second screening RipAB, RipAC, RipAI, RipAX1, RipAY, RipBM, RipC1, RipE1, RipH1, RipN, RipR, RipS4, RipU, RipX and RipZ were screened but no targets were found, and RipAN and RipM could not be screened because of recalcitrant problems with yeast transformation. For *Xcc* strain 8004: AvrXccA2, HpaA, HrpW, XopAN, XopN and XopQ were screened but no targets were found, and AvrBs2, XopAH, XopAL2, XopD and XopE2 could not be screened because they showed autoactivation in yeast.

**S2 Fig. Overlap of *Ath* targets of effector proteins from *Hpa*, *Psy*, *Gor*, *Rps* and *Xcc*.**

Venn diagrams showing the overlap among *Ath* targets found in the 8,000-*Ath*-cDNA collection (8K space) of effector proteins from *Hpa*, *Psy*, *Gor*, *Rps* and *Xcc* at the kingdom (A) and species level (B). The total number of effector targets for each kingdom/species is indicated in brackets.

**S3 Fig. Interspecific convergence of *Psy* and *Xcc* effector proteins.**

Number of *Ath* targets in the 8K space of effectors from *Psy* and *Xcc* and *Rps* strain found in 10,000 degree-preserving simulations (grey) versus the observed number (red arrow).

**S4 Fig. Effector degree distribution for *Ath* effector targets.**

Effector degree (i.e., number of effectors that interact with an *Ath* protein) distribution among the 564 identified *Ath* effector targets (A), according to the origin the data: published large-scale screenings in light green, manual curation of literature in mid-green and this study in dark grey or (B), according to the kingdom of the corresponding effector pathogen: Bacteria in light blue, Chromista in dark blue, Fungi in light orange and Animalia in dark orange.

**S5 Fig. *Ath* degree and betweenness centrality of different groups of *Ath* effector targets.**

Cumulative distribution of *Ath* degree (B and D) and betweenness centrality (A, C and E) for *Ath* proteins targeted (orange) or not (purple) by effectors (B), multi-pathogen (green) and pathogen-specific (pink) effector targets (B and C) and effector hubs (red) and single effector targets (blue) (D and E). The significance of the differences were evaluated by one-tailed Wilcoxon signed-rank test. The illustration in the upper right corner of each graph represents each compared group: effectors are represented by squares, *Ath* proteins by circles, numbers represent different pathogens species and the color code matches the respective cumulative distribution graph. The estimation of the area under the curve of each distribution is compiled in Table 2.

**S6 Fig. *Ath* and effector degree of effector targets.**

(A) Scatterplot of *Ath* degree versus effector degree of all *Ath* effector targets. Squared in a grey dashed line is the close-up area represented in (B).

**S7 Fig. Degrees and betweenness centrality of bacterial core and non-core T3Es and their targets.**

Cumulative distribution of *Ath* degree (A and D), effector degree (C) and betweenness centrality (B and D) for bacterial core T3Es (yellow) and other bacterial T3Es (cyan) (A and B) and their targets (brown and blue respectively) (C-E). The significance of the differences were evaluated by one-tailed Wilcoxon signed-rank test. The illustration in the upper right corner of each graph represents each compared group: bacterial T3Es are represented by squares, *Ath* proteins by circles and stars represents bacterial core T3Es. The estimation of the area under the curve of each distribution is compiled in Table 2.

S1 Table. List of *Rps* and *Xcc* effector-*Ath* protein interactions detected experimentally in this study and composition of the *Ath*-cDNA screening libraries.

S2 Table. List of manually curated *Ath*-effector protein interactions from the literature.

S3 Table. List of effector hubs and single effector targets identified.

S4 Table. List of protein interactions constituting the *Ath*-effector interactomic network.

S5 Table. List of pENTRY for T3E genes from *Rps* and *Xcc*.

